# A fluorescent sex-sorting technique for insects with the demonstration in *Drosophila melanogaster*

**DOI:** 10.1101/2023.08.11.553026

**Authors:** Junru Liu, Danny Rayes, Omar S. Akbari

**Author notes:** To whom correspondence should be addressed: Omar S. Akbari, Ph.D., School of Biological Sciences, Department of Cell and Developmental Biology, University of California, San Diego, La Jolla, CA 92093, USA.

## Abstract

Recent advances in insect genetic engineering offer alternative genetic biocontrol solutions to control populations of pests and disease vectors. While success has been achieved, sex-sorting remains problematic for scaling many genetic biocontrol interventions. Here we describe the development of a sex-sorting technique for female and male selection with a proof-of-concept in *D. melanogaster* termed SEPARATOR (<u>S</u>exing <u>E</u>lement <u>P</u>roduced by <u>A</u>lternative <u>R</u>NA-splicing of <u>A</u> <u>T</u>ransgenic <u>O</u>bservable <u>R</u>eporter). This approach utilizes dominant fluorescent proteins and differentially spliced introns to ensure sex-specific expression. The system has the potential for adaptability to various insect species and application for high-throughput insect sex-sorting.

## Introduction

Genetic biocontrol methods are highly effective and sustainable alternatives to traditional insecticide-based approaches for suppressing pest and vector populations ^1,2^. Sterile insect technique (SIT), for example, is a genetic biocontrol technology that has successfully eradicated screwworms in the United States ^3,4^. This technology suppresses populations through the frequent releases of sterile males, as female insects are often responsible for destroying agricultural resources or pathogen transmission ^5^. While SIT has proven to be effective in some species, it requires the release of only sterile males, necessitating the development of technologies to efficiently sort the sexes. This indeed remains a grand challenge for many genetic biocontrol technologies and what is needed is a technology that can be rapidly developed in many species to enable rapid and accurate sex sorting.

Early sex sorting was a labor-intensive process that involved manually sorting individuals by their sexual dimorphisms, severely limiting throughput and scalability. More recent approaches utilize machine learning to automate sex sorting ^6,7^. However, these still rely on natural sexual dimorphisms, making them challenging to develop to other species.

Existing genetic approaches for sex-sorting insects also have limitations that impact their scalability. The classic Genetic Sex Separation (GSS) method involves irradiation-induced translocation of a conditional lethal gene, such as the gene conferring resistance to the insecticide dieldrin (Rdl), to the Y chromosome ^8^. As a result, only males carrying the translocated gene on their Y chromosome can survive when exposed to the insecticide. This GSS method has succeeded in certain insect species, but overall this method is difficult to develop and prone to breakage via meiotic recombination.

A scalable sex-sorting method is essential to ensure cost-effective and scalable genetic technologies are available for the control of a variety of insect species. Incorporating fluorescent sex-specific markers facilitates the visual identification and separation of males and females with a Complex Parametric Analyzer and Sorter (COPAS, Union Biometrica) instrument. The COPAS has been used to sort *D. melanogaster* embryos and larvae based on fluorescence as well as transgenic *Aedes aegypti* and *Anopheles gambiae* larvae ^9,10^. The COPAS instrument was used to fully automate the sorting process, achieving the efficiency, speed, and accuracy needed to scale genetic technologies ^10^.

Here we describe the proof of principle for a sex-sorting technique termed SEPARATOR (<u>S</u>exing <u>E</u>lement <u>P</u>roduced by <u>A</u>lternative <u>R</u>NA-splicing of <u>A</u> <u>T</u>ransgenic <u>O</u>bservable <u>R</u>eporter) that utilizes fluorescent proteins and sex-specific introns of the sex-determination gene *transformer (tra)* to ensure sex-specific expression. Female-specific expression is achieved by disrupting the coding sequences (CDS) of a fluorescent reporter with female-specific *transformer* introns (*traF*). These female-specific introns are only spliced in females, resulting in the restoration of gene function and female-specific fluorescence. In this project, *traF* from four separate species were tested, including *Drosophila melanogaster*, *Drosophila suzukii, Ceratitis capitata*, and *Anastrepha ludens*. When *traF* from *D. melanogaster, D. suzukii*, and *C. capitata* is inserted into the CDS of fluorescence protein, we achieved 100% female-specific fluorescence. Notably, *C. capitata traF* permits the selection of females as early as L1 instar larval stage. However, *A. ludens traF* resulted in dsRed expression in both sexes in *D. melanogaster* flies, indicating that *A. ludens traF* is not sex-specific when transcribed in *D. melanogaster*. This SEPARATOR technique is a valuable method for insect sex-sorting as it (1) exploits highly conserved sex-specific splicing mechanisms, making it widely transferable to different insect species; (2) allows 100% positive selection for either females or males based on fluorescence instead of morphological differences, providing better accuracy; (3) permits sex-sorting during different life stages and as early as L1 instar stage; (4) can be combined with existing genetic control methods and a COPAS machine for precise high-throughput screening.

## Results

### Development of female-specific expression of the fluorescent protein

To engineer female-specific expression of a reporter for positive selection of females, we exploited the sex-specific alternative splicing of a conserved sex-determination gene. In *D. melanogaster*, the *transformer (tra)* intron between exons 1 and 2 is spliced out in females, resulting in a functional *tra* protein. In males, alternative *tra* splicing results in a premature stop codon that terminates the *tra* protein

^11^. This female-specific alternative splicing mechanism occurs not only in *Drosophila* but also in *Ceratitis* and *Anastrepha*, suggesting that it is highly conserved in Dipterans (**Fig. 1a, Fig. s1 & s2**) ^12^. Consequently, the *traF* from *D. melanogaster, D. suzukii, C. capitata*, or *A. ludens* were inserted into the fluorescent protein coding sequences to test for female-specific fluorescent protein expression (**Fig. 1b**). We generated two sets of dual fluorescent marker constructs encoding a fluorescent marker for both sexes and a female-specific fluorescent marker (**Fig. 1c**). The constructs were cloned into a plasmid containing an attP recombination site and a *piggyBac* (PB) transposable element. Set 1 constructs have an eGFP fluorescence expressed under a ubiquitous promoter *Hr5Ie1* (*Hr5Ie1*-eGFP) as the selectable marker for the transgene. To promote constitutive expression of female-specific fluorescent proteins, we used another ubiquitous promoter *Opie2* to express dsRed and inserted *traF* immediately downstream of the ATG translational start codon of dsRed (*Opie2*-ATG-*traF*-dsRed). Constructs in set 2 have the opposite marker configuration, with *traF* inserted downstream of the ATG translational start codon of eGFP under promoter *Hr5Ie1* (*Hr5Ie1*-ATG-*traF*-eGFP) for the female-specific fluorescent expression and *Opie2*-dsRed as the selectable marker for the transgene. In total, nine constructs were created: a control construct (795G) and eight experimental constructs, with four constructs in each set (**Fig. 1d**).

**Fig. 1.**
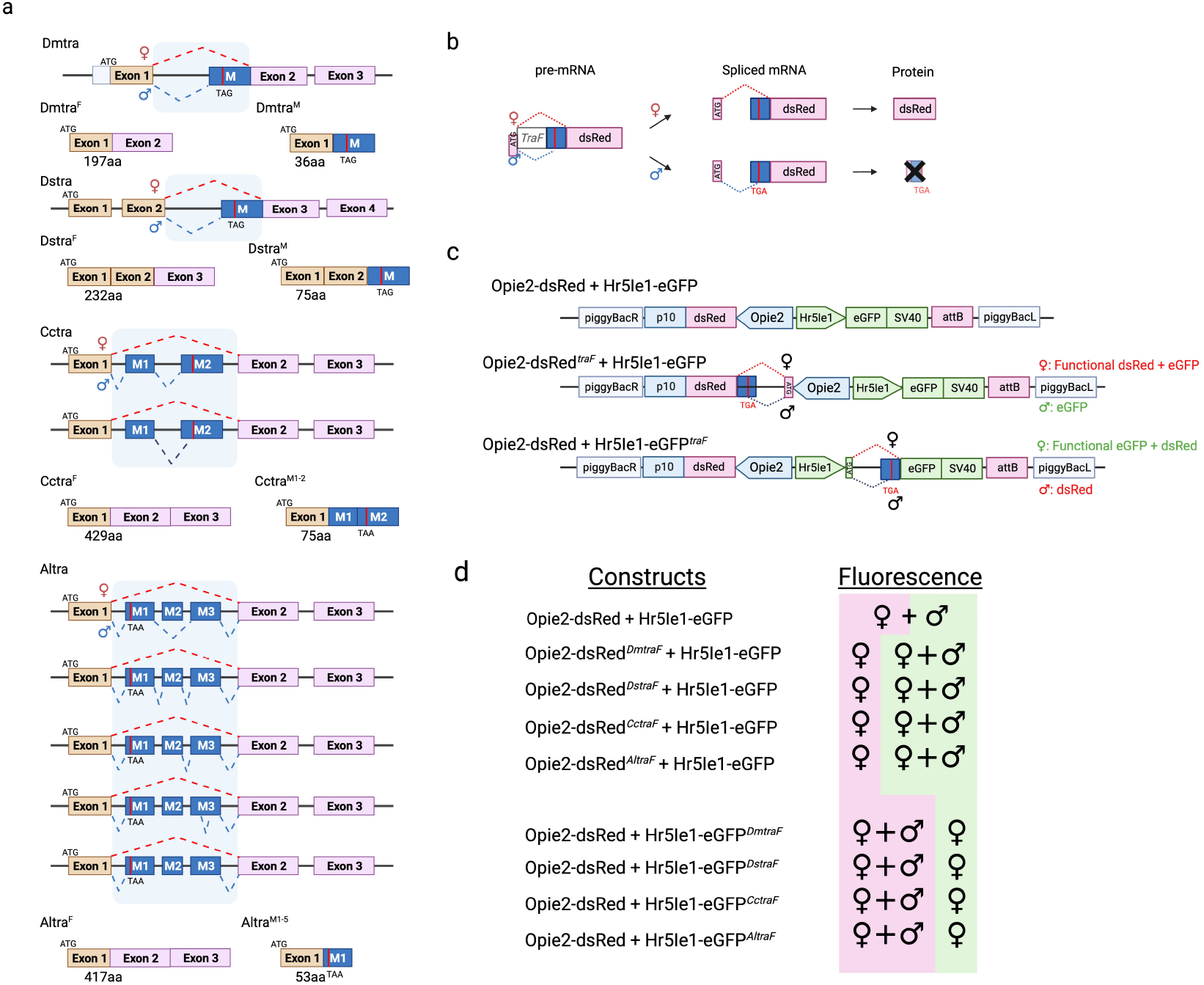
Sex-sorter cassette in *Drosophila*. (a) Sex-specific alternative splicing and the resulting protein of the *transformer (tra)* in *D. melanogaster, D. suzukii, C. capitata*, and *A. ludens*. (b) Splicing of the female-specific *transformer* (*TraF*) intron should result in functional dsRed protein in females but not in males. (c) Schematic of the sex-sorter constructs engineered and tested in the study. *TraF* introns from *D. melanogaster, D. suzukii, C. capitata*, and *A. ludens* are inserted into the coding sequence of either dsRed or eGFP after the ATG translational start codon. (d) Fluorescence expression of females and males carrying the respective constructs.

### *DmtraF, DstraF*, and *CctraF* resulted in female-specific fluorescence

The transgene integration site can impact gene expression, so we opted to integrate all nine constructs into the same site through phiC31 *attP* integration on the second chromosome (BDSC #25709). Nine homozygous transgenic strains were established. Six constructs that harbor *DmtraF, DstraF*, and *CctraF* resulted in female-specific fluorescence (795H,I,J,L,M, and N, **Fig. 2**). These results indicate that inserting the *traF* into the coding sequence of the fluorescence proteins can result in female-specific fluorescence. However, for constructs harboring *AltraF* (795K and O), both females and males exhibited the intended female-specific fluorescence (**Fig. 2**). This result suggests that *AltraF* is spliced out in both females and males rather than in a female-specific manner.

**Fig. 2.**
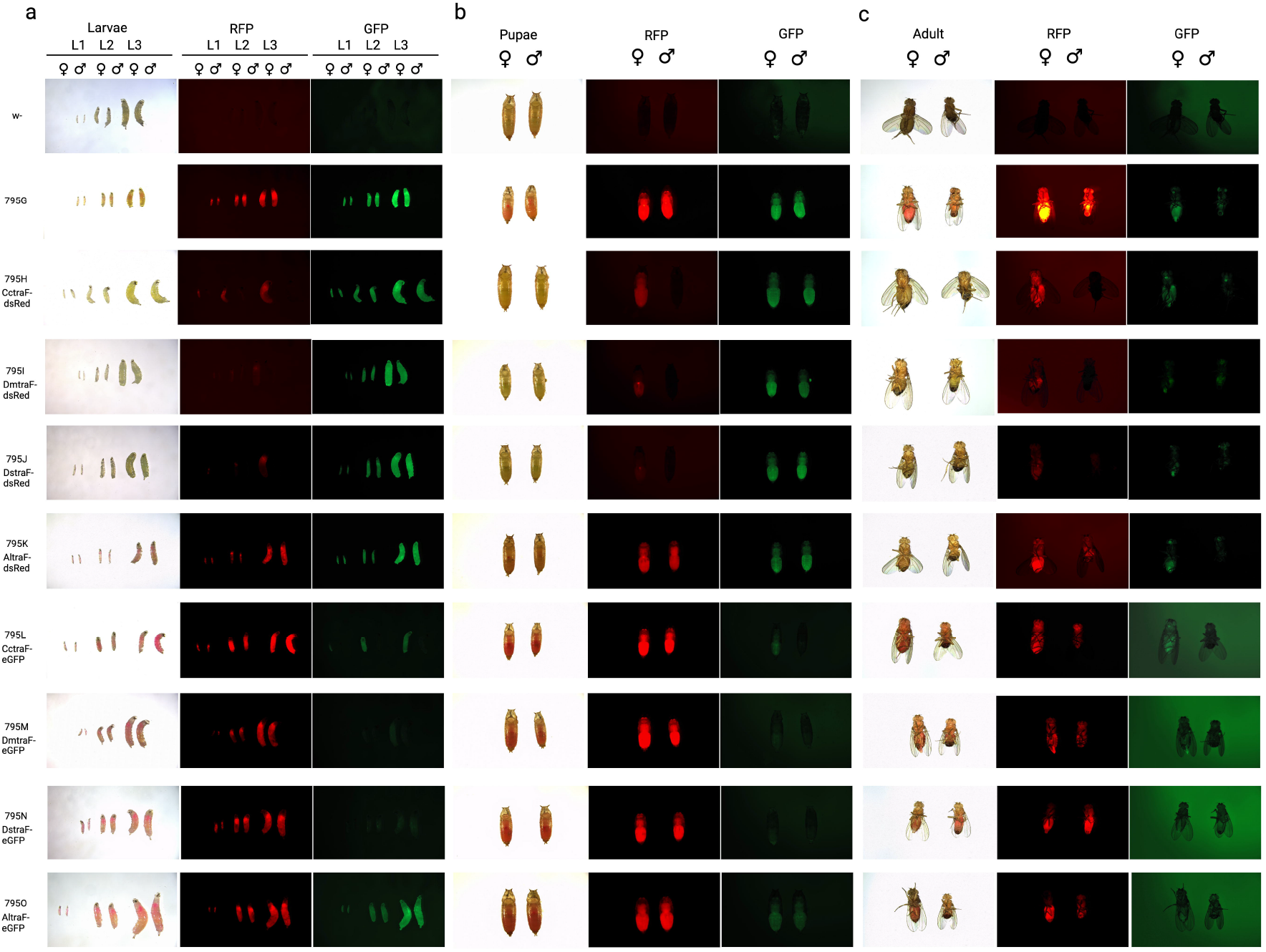
Expression of Opie2-*TraF*-dsRed or Hr5ie1-*TraF*-eGFP transgenes in *D. melanogaster* can be observed in different developmental stages (a) L1-L3 larval stage, (b) pupal stage, (c) adult under white light, RFP and GFP filters.

To validate the alternative splicing variants, adult flies were collected for RT-PCR analysis to obtain the fluorescent protein transcripts. Primers were designed to anneal to the 5’ UTR region at the 3’end of the *Opie2* promoter, and the 3’ end of the dsRed sequence (**Supplementary Table 1**). Multiple bands were obtained from the RT-PCR samples for *CctraF* males, and both sexes of *DmtraF* and *DstraF* (**Fig. s3a &b**). Sequencing of these bands indicates that *CctraF, DmtraF*, and *DstraF* resulted in functional dsRed expression explicitly in females, while *AltraF* had dsRed expression in both males and females (**Fig. s3c**). The molecular results obtained from RT-PCR analysis were consistent with our observations in the flies, confirming that *CctraF, DmtraF*, and *DstraF* exhibited female-specific splicing in *D. melanogaster*. This outcome demonstrates the feasibility of this fluorescent sex-sorting approach, as these female-specific splicing events allow for the positive selection of either sex.

### Confirmation of sex-specific fluorescence at multiple developmental stages

Next, we evaluated the intensity and sex specificity of the fluorescence over multiple life stages. The six constructs (795H, I, J, L, M, and N) that exhibited female-specific fluorescent expression were evaluated. Female-specific fluorescence was observed as early as in the first instar larvae (L1) life stage in both *CctraF* transgenic lines: 795H and 795L (**Fig 2a, Fig 3, Supplymentary Table 2**). Female-specific fluorescence was also observed in the third instar larvae (L3) of *DmtraF* 795I and *CctraF* 795L, and the pupal stage for *DmtraF* 795M and *DstraF* 795N. Despite the identical introns in the *DmtraF* 795I and *DmtraF* 795M and the *CctraF* 795L and *DstraF* 795N strains, female-specific fluorescence was detected earlier in strains with the female-specific dsRed marker. This result is presumably due to the deeper tissue penetrance and lower auto-fluorescence of red fluorescent protein (RFP) ^13^. Notably, the intensity of the female-specific fluorescence varies among introns. The *CctraF* exhibits the highest brightness, followed in order of brightness by *DmtraF* and *DstraF* (**Fig. 2**). This is unexpected as *CctraF* is an exogenous/non-native intron for *D. melanogaster*, potentially hindering successful intron recognition and splicing efficiency.

**Fig. 3.**
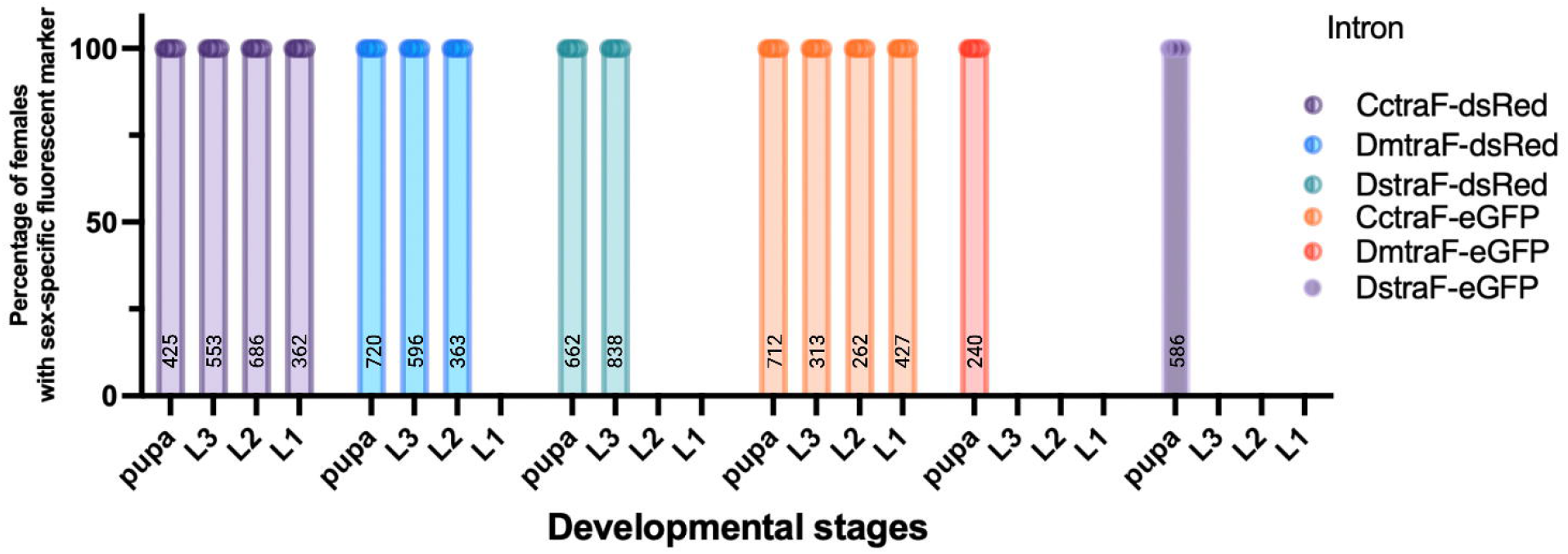
Female selection efficiency at different life stages in *D. melanogaster* for all six sex-sorter cassettes that give female-specific fluorescence and the numbers of scored flies are indicated for each bar.

### Assessing the fitness of the sex-sorting strains

Strain fitness is essential for scalability. Fluorescent proteins have documented fitness costs to genetically engineered organisms, but we expected that including *traF* in their coding sequences would minimally affect the fitness of the sex-sorting strain. We, therefore, compared the egg hatching and larval to adult survival rate of all eight homozygous sex-sorting strains to a control strain (795G) containing fluorescent reporters lacking *traF* introns. Our finding indicates that there is a significantly higher hatching rate in flies harboring *Opie2-ATG-traF-dsRed* constructs when compared to 795G control (**Fig. 4**). This effect could possibly be attributed to the fitness cost associated with dsRed functioning as a tetramer ^14^. With the inclusion of *traF* introns, the expression of dsRed occurs at a reduced level. As a consequence, the fitness cost is diminished, which, in turn, leads to a higher hatching rate. We observed lower larvae to adult survival only in the CctraF-dsRed 795H strain (p< 0.05, Student’s t-test with equal variance, **Fig. 4)**. These results suggest that the *traF* intron does not impose substantial fitness costs on the strain, making them suitable for potential large-scale insect population control projects.

**Fig. 4.**
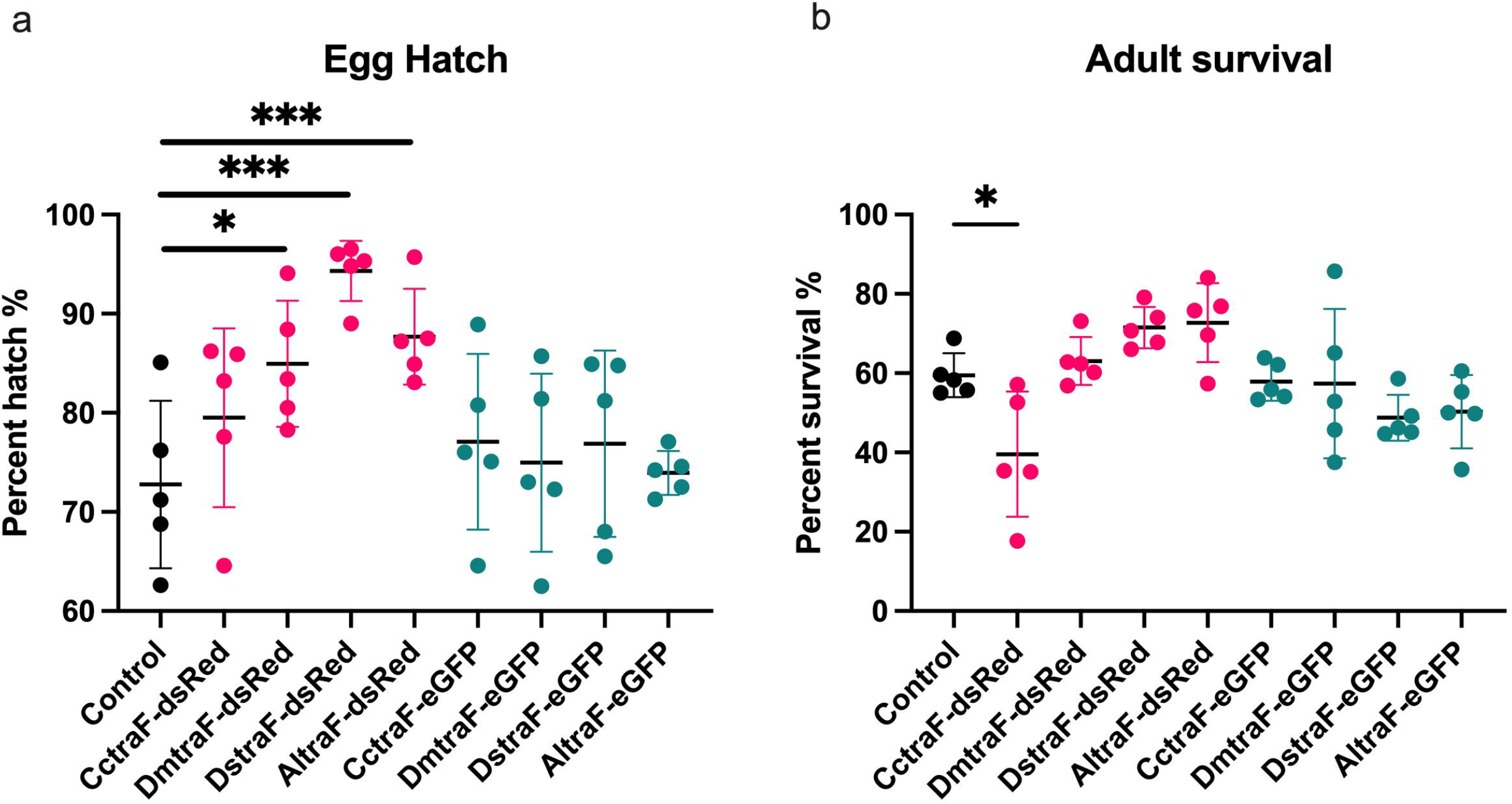
Fitness cost of all eight sex-sorting cassettes was accessed through two parameters: (a) egg-hatching rate and (b) survival rate to adulthood. Fitness cost was observed in *CctraF-*dsRed strain in the parameter of survival rate to adulthood (*p<0.05, ***p<0.001, Student’s t-test with equal variance.)

## Discussion

In this study, we engineered a novel sex-sorting method, SEPARATOR, that integrates a female-specific intron into the coding sequencing of a fluorescent reporter to generate an efficient female-specific marker. This innovative technique uses splicing mechanisms conserved across dipteran species, and has even been successfully engineered in other dipteran species ^15^. We examined the splicing patterns of four *traF* introns, *DmtraF, DstraF, CctraF*, and *AltraF*, and found that three of them *(DmtraF, DstraF*, and *CctraF)* had 100% female-specific splicing. Among the six constructs tested for female-specific fluorescence expression in *D. melanogaster*, those carrying the *CctraF* introns (795H and 795L) exhibited expression at the earliest life stage and the brightest expression. These strains can be used for sex-sorting as early as the L1 stage and throughout the entire life cycle. This observation aligns with previous studies demonstrating the functional conservation of *CctraF* and its applicability in various insect species ^16–18^.

It is worth noting the significant size difference between the *traF* introns in Drosophilidae species (*DmtraF* and *DstraF*) and Tephritidae species (*CctraF* and *AltraF*). The *traF* introns in Drosophilidae species are approximately 200 bp in length, whereas in Tephritidae species, they are approximately ten times longer. Although *AltraF* is not female-specifically spliced, it is spliced out entirely in both females and males, thus resulting in functional fluorescent expression in both sexes. This result could be attributed to species differences in splicing signals as *AltraF* is a foreign intron in *D. melanogaster*. These size differences may also impact intron splicing efficiency and, consequently, the dual sex expression of the *AltraF* fluorescent reporter. Larger introns, for example, possess more heterogeneous nuclear ribonucleoproteins (hnRNP) and serine/arginine-rich (SR) proteins recognition sites, leading to higher splicing efficiency ^19,20^. However, further testing in Tephritidae species like *C. capitata* or *A. ludens* may better elucidate the capabilities of *AltraF* intron for SEPARATOR systems.

The *CctraF* introns, on the other hand, generate efficient and easily screenable female-specific fluorescent phenotypes. This efficiency may be due to the simplified splicing of *CctraF*, which does not depend on the presence of the *sex-lethal (sxl)* protein required in *Drosophila* for *tra* transcript splicing ^16,21–24^. In *C. capitata*, the *tra* gene is autoregulated and continuously expressed from the embryo to adulthood ^16^, allowing for the consistent and stable splicing of *CctraF*. Therefore, *CctraF* may be a versatile tool for sex-specific expression in a wide range of insect species.

SEPARATOR has numerous advantages over traditional sexing methods. These include its potential multiple species adaptability, positive selection based on dominant fluorescence for both sexes, sex differentiation during early stages of development, genetic stability, and minimal impact on fitness. All of these characteristics improve the potential scalability and timing of sex sorting. The fluorescent sex-sorting cassettes were engineered to be highly transferable to numerous insect species. They utilize a PB transposable element, which has high transgenesis efficiency in many insect species ^25^. The two constitutive baculovirus promoters for fluorescent protein expression, *Opie2* and *Hr5Ie1*, have been shown to facilitate high expression in various insects throughout development ^26–28^. Furthermore, dsRed and eGFP are the two most commonly used fluorescent proteins and can penetrate most insect tissues ^29^. Female-specific alternative splicing of *tra* intron is also highly conserved across insect species ^24,30^.

The SEPARATOR technology enables the positive selection of females, or males, in all post-embryonic developmental stages. Sex sorting at earlier developmental stages would be advantageous for large-scale insect population control projects, as it would extend the sex sorting timeline, facilitate the release of earlier life stages, and may reduce the costs of large-scale releases ^31^. Our approach simplifies manual sex sorting, but the primary benefit is to large-scale sex-sorting applications. SEPARATOR can be combined with existing genetic control methods and a COPAS machine for precise and high-throughput screening. Fluorescence can also serve as a reliable indicator of undesired genetic events such as loss-of-function (LOF), gain-of-function (GOF), or genetic exchange such as chromosomal translocation and recombination that would result in loss of fluorescence. The GSS used for many current SIT applications rely on reciprocal chromosomal translocations between the Y-chromosome and a region of the autosome containing a selectable marker. These GSS methods have been designed for male-only release programs, where females possess distinct phenotypes or markers that facilitate easy identification and separation ^32^. However, these GSS methods are prone to genetic instability and can be disrupted by meiotic recombination or chromosomal rearrangements. Studies on the multi-generation of GSS lines have revealed that chromosomal recombination can occur at a frequency of approximately 0.07%, leading to the breakdown of the sexing system ^33^. To enhance stability, one potential solution is to generate new translocations with a breakpoint located in close proximity to the sexing genes ^33^.

Although the main emphasis of this work has sex-sorting applications, SEPARATOR can be used to study fly development. Differentiation between female and male adult flies is easily accomplished under the microscope, but accurate larvae sex-sorting can be challenging, particularly when they are embedded within the food medium. The major distinguishing factor between female and male larvae is the presence or absence of gonads, which is not easily visible ^34^. With the *CctraF* 795H or 795L sex-sorter cassettes, the fluorescent protein markers can be easily observed as early as first instar larvae, even in the food medium and with minimal autofluorescence. The ability to accurately sex-sort larvae at early developmental stages provides a valuable tool for tracking the sex of flies and could potentially facilitate the study of sex-specific developmental processes in flies.

## Materials and Methods

### Molecular cloning

All genetic constructs were produced utilizing the Gibson enzymatic assembly. The construct 795G was created using a pre-existing plasmid containing piggyBac, attB-docking sites, and an *Opie2* promoter regulating dsRed. This plasmid was subsequently linearized with XhoI and NotI enzymes. The *Hr5Ie1* promoter, along with eGFP were cloned into the linearized plasmid to make 795G, which serves as the control plasmid. To generate female-specific dsRed (795H-K), the plasmid 795G was linearized with AvrII and BamHI to allow insertion of introns into dsRed. Alternatively, to generate female-specific eGFP (795L-O), 795G was linearized using MluI and BsrGI to insert introns into eGFP. The *traF* introns from *D. melanogaster, D. suzukii, C. capitata*, or *A. ludens* were amplified from their respective genomic DNA using the primers listed in the Supplementary Table 1.

### Reverse transcription PCR (RT-PCR) of the female-specific splicing transcripts

To access the splicing transcripts of four *traF* introns, we screened for female- and male-specific dsRed mRNA. Total RNA of ten virgin females or males from w-, 795G, H, I, J, and K were extracted using the miRNeasy Tissue/Cells Advanced Kits (Qiagen). DNase treatment is done using the TURBO™ DNA-free (Invitrogen), and followed by the cDNA synthesis using the RevertAid First Strand cDNA Synthesis Kit (Thermo Scientific™). The genomic DNA (gDNA) was amplified using primers 795.s2F and 795.s2R, and the cDNA was amplified using primers 795.s3F and 795.s1R (Supplementary Table 1). The gDNA samples were run on 1% TAE agarose gel, and the cDNA samples were run on a 2% TAE agarose gel.

### Rearing and fly transgenesis

Transgenic flies were maintained under standard conditions at 25ºC with a 12H/12H light/dark cycle and fed on the Old Bloomington Molasses Recipe. Embryonic injections were performed in the lab following the standard injection protocol. Plasmids diluted to 300-350ng/µL in water were inserted at P{CaryP}attP40 on the 2nd chromosome (Bloomington #25709). Recovered transgenic lines were balanced on the 2nd chromosome using a single chromosome balancer line w1118; CyO/sna[Sco]. Multiple independent lines were obtained for each plasmid and tested for sex-specific fluorescence. We used homozygous transgenic lines containing two copies of the transgene to assess the sex-sorting efficiency. Sex-sorting lines with *CctraF* introns 795H and 795L are deposited at the Bloomington Drosophila Stock Center (BDSC# pending).

### Genetics and sex selection

To assess the fluorescent sex selection efficiency, we crossed ten virgin females to ten males in a fly vial. The parental flies were flipped into a fresh vial every 12hrs, and the numbers of the laid embryos were scored. After hatching, the larvae or pupa were scored and transferred to different vials based on their fluorescent markers. The sex and the fluorescent markers of the adult offsprings were recorded after eclosion. Flies were scored using a Leica M165FC fluorescent stereomicroscope. Images were taken using a View4K camera. Each genetic cross was set up five times using different parental flies.

### Fitness estimation

The fitness of the sex-sorting strains is assessed based on two parameters: the rate of egg-hatching (from embryos to larvae) and the rate of adult survival (from larvae to adult). To evaluate the egg-hatching rate, flies are allowed to lay embryos in fly vials for a duration of 24 hrs, and the number of eggs laid in each vial is recorded. After 24 hrs of egg laying, the number of larvae is recorded. To assess the adult survival rate, the number of both female and male adult flies that successfully eclosed is recorded.

### Statistical analysis

Statistical analysis was performed in Prism9 by GraphPad Software, LLC. Three to five biological replicates were used to generate statistical means for comparisons.

## Data availability

Complete sequence maps and plasmids are deposited at Addgene.org (#205481-205489). Transgenic lines 795H and 795L have been made available for order from Bloomington Drosophila stock center. The other transgenic lines are available upon request to O.S.A.

## Acknowledgements

This work was funded by the United States Department of Agriculture (USDA) - Animal and Plant Health Inspection Service (APHIS) - Plant Protection and Quarantine (PPQ) (AP19PPQS&T00C237 and AP22PPQS&T00C188) and funding from an NIH award (R01AI151004) awarded to O.S.A. The views, opinions, and/or findings expressed are those of the authors and should not be interpreted as representing the official views or policies of the U.S. government. Figures were created with BioRender.com.

## Author contributions

O.S.A conceived and designed the experiments. J.L and D.R performed molecular and genetic experiments. All authors contributed to the writing, analyzed the data, and approved the final manuscript.

## Figure legends

**Supplementary Figure 1.**
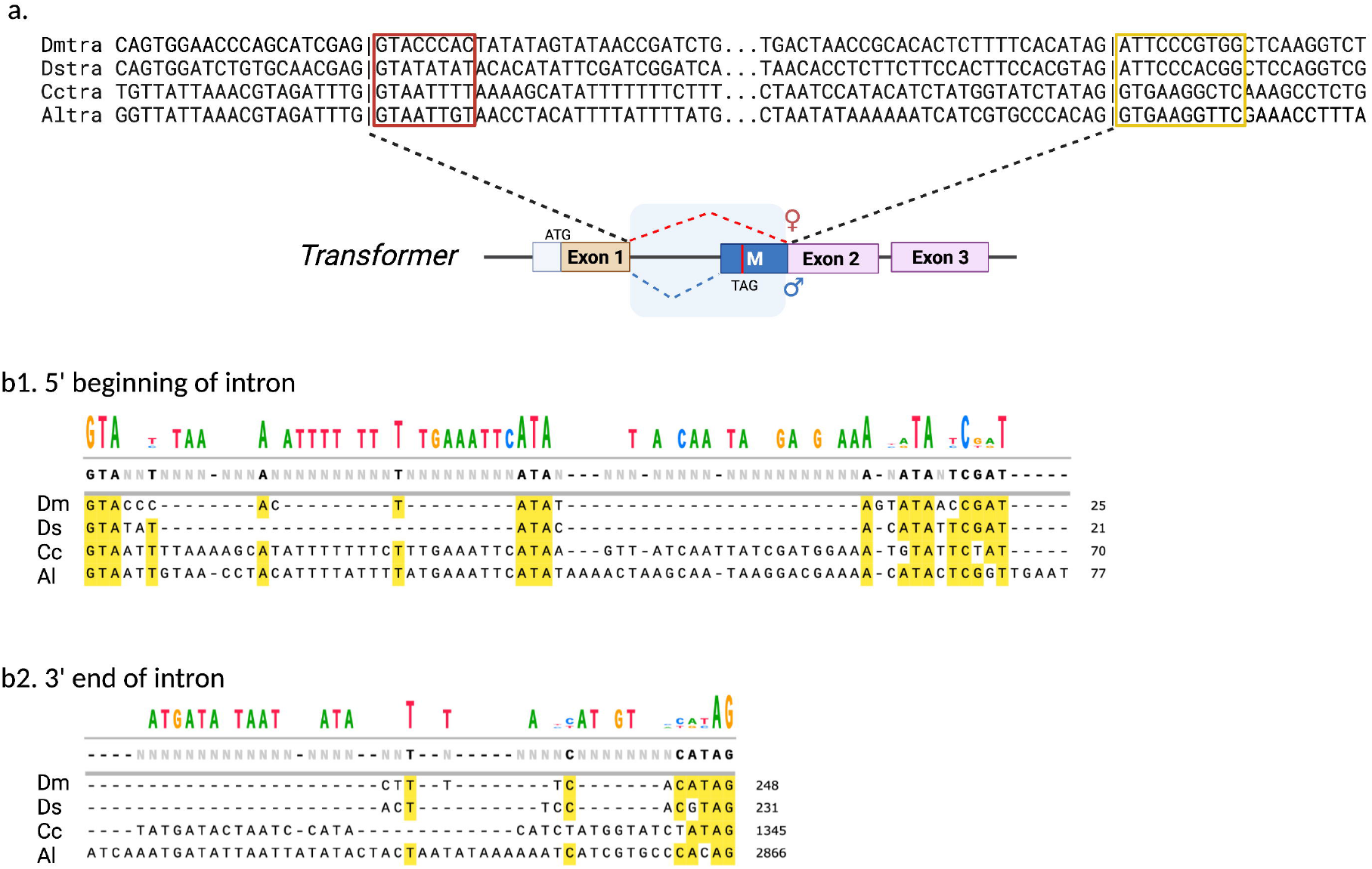
(a) Comparison of the *transformer* female-specific intron splice donor (red square) and acceptor sites (yellow square) from *D. melanogaster, D. suzukii, C. capitata*, and *A. ludens*. (b) Alignment of the sequences at the 5’ beginning (b1) and the 3’ end (b2) of the intron.

**Supplementary Figure 2.**
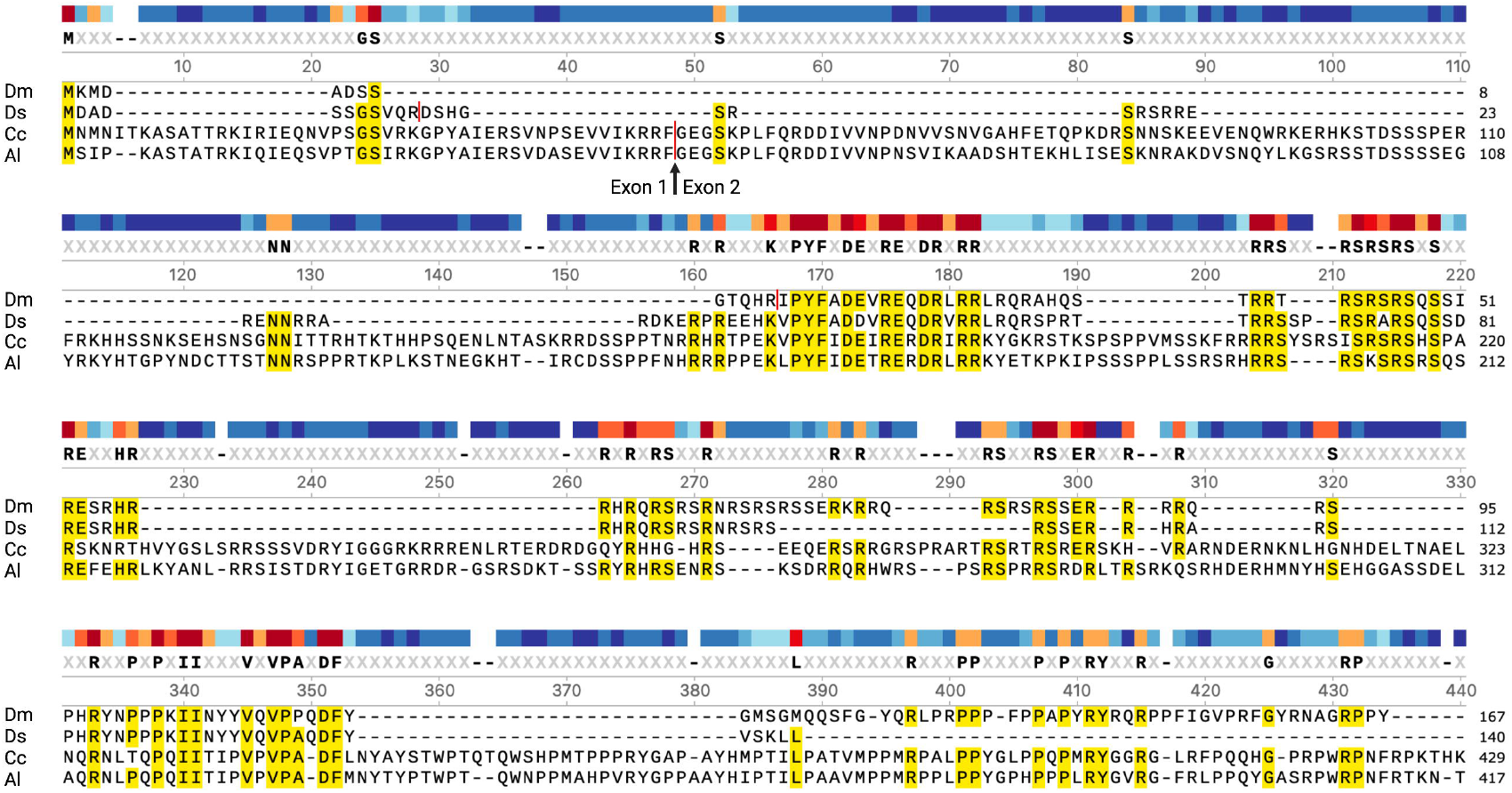
Protein alignment of the *transformer* protein in *D. melanogaster, D. suzukii, C. capitata*, and *A. ludens*. Conserved amino acids among four species are highlighted in yellow.

**Supplementary Figure 3.**
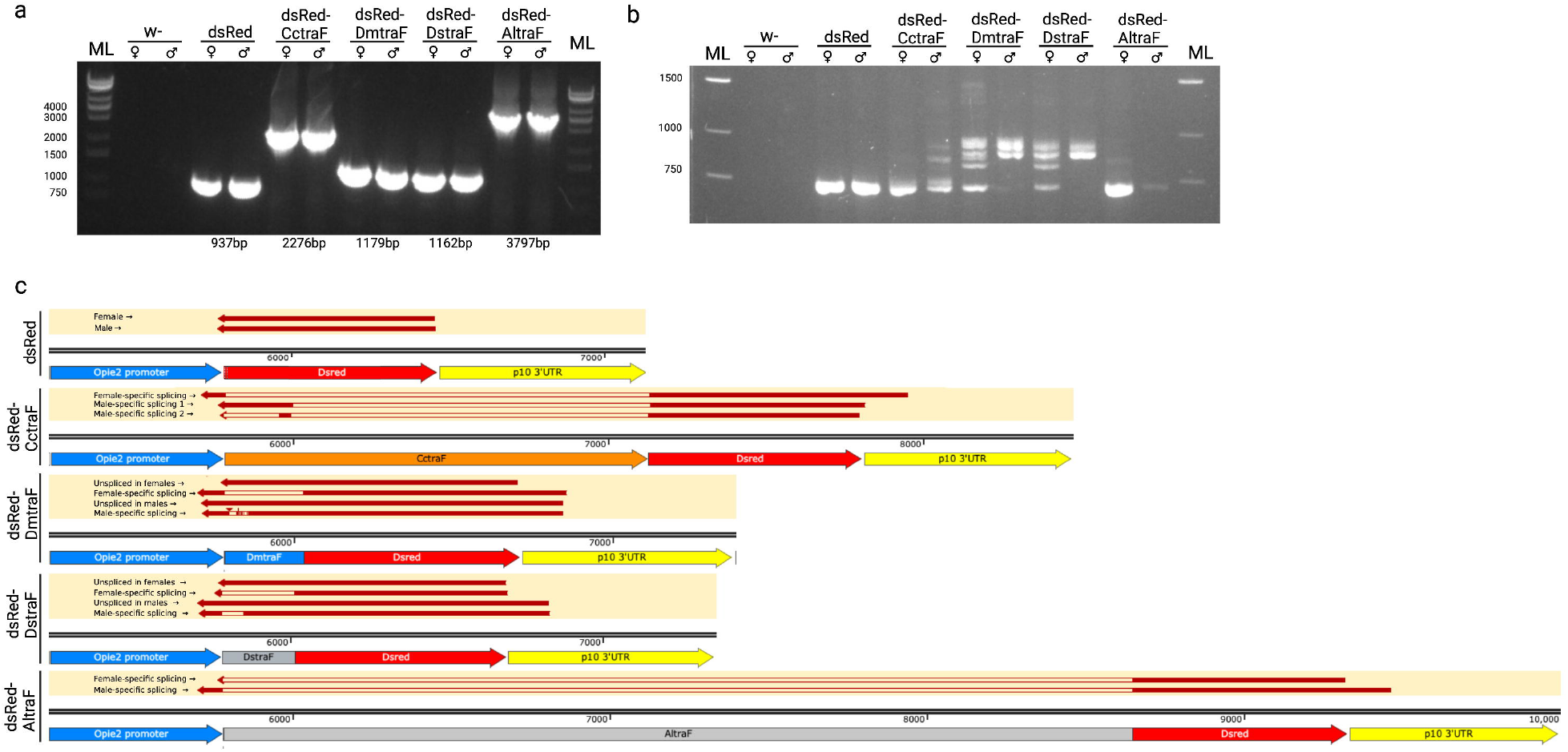
The *traF* introns splicing patterns in *D. melanogaster*. Gel electrophoresis images show (a) the genomic DNA PCR for dsRed-*traF* (b) the cDNA for the dsRed-*traF*. ML: molecular ladder. (c) sequencing results of the cDNA from each band.

**Supplementary Table 1:** Primer sequences used in this study.

**Supplementary Table 2:** Sex-specific fluorescence sorting at multiple developmental stages.

