## Supplementary Table 1 for "A fluorescent sex-sorting technique for insects with the demonstration in *Drosophila melanogaster*"

Supplementary Table 1: Primer sequences used in this study.

| | | Primer name | Primer seequence (5'-3') |
|---|---|---|---|
| genotypeing introns | DstraF | 1122.S1F | cggctaggttaaacgagtagtgtccacctc |
| | | 1122.S2R | GTAATTTACTAACGTAGAAATCCTGTGCTGGCAC |
| | CctraF | 825A | AAAAATAAGGTCAGCAGCCATGAC |
| | | 825B | GTCATGGCTGCTGACCTTATTTTT |
| | | 825C | AGACATTCCAAATTCAAGTTAACAAATTAAT |
| | | 825D | ATTAATTTGTTAACTTGAATTTGGAATGTCT |
| | AltraF | 1168-Tra-gDNA-R1 | CTTGCTCTGGAGTCCGTTAAA |
| | | 1168-Tra-gDNA-F2 | CCTGAGCAGATAACCCTCTTTC |
| | | 1168-Tra-gDNA-F3 | GGTCGTTTCGGTAACGTAGAA |
| | | 1168-Tra-gDNA-R2 | GTCATCAGTTGTCGGCATTTAC |
| Cloning primers | | 795G2.c1R | ccgcaacctgtctctggtgGGATCCatggtgcgctcctccaagaacgtcatcaag |
| | | 795K | acgccatccaaccgccgccgcaacctgtctctggtgATGgtaattttaaaagcatatttttttc |
| | | 795I | acgccatccaaccgccgccgcaacctgtctctggtgATGgtaattttaaaagcatatttttttc |
| | | 1122J.c1F | aattcaacgcacacttattacgtgaggtaccgcgcccatactcggtggcctccccac |
| | | 795L.c1F | ggccgactgttttcgtatccgctcaccaaacgcgtttttgcattaacattgtatgtcggc |
| | | 795L.c2R | tcaaagaaaaaaatatgcttttaaaattacCATtttgTATTgtcacttggttgttcacga |
| | | 795L.c3F | tcgtgaacaaccaagtgacAATAcaaaATGgtaattttaaaagcatattttttttcttt |
| | | 795L.c4R | ACCCCGGTGAACAGCTCCTCGCCCTTGCTctatagataccatagatgtatggattagtat |
| | | 795L.c5F | atactaatccatacatctatggtatctatagAGCAAGGGCGAGGAGCTGTTCACCGGGGT |
| | | 795M.c2R | ATCAGATCGGTTATACTATATAGTGGGTACCATtttgTATTgtcacttggttgttcacga |
| | | 795M.c3F | tcgtgaacaaccaagtgacAATAcaaaATGGTACCCACTATATAGTATAACCGATCTGAT |
| | | 795M.c4R | CACCCCGGTGAACAGCTCCTCGCCCTTGCTCTATGTGAAAAGAGTGTGCGGTTAGTCAAT |
| | | 795M.c5F | ATTGACTAACCGCACACTCTTTTCACATAGAGCAAGGGCGAGGAGCTGTTCACCGGGGTG |
| | | 795N.c2R | tctgatccgatcgaatatgtgtatatatacCATtttgTATTgtcacttggttgttcacga |
| | | 795N.c3F | tcgtgaacaaccaagtgacAATAcaaaATGgtatatatacacatattcgatcggatcaga |
| | | 795N.c4R | CCCCGGTGAACAGCTCCTCGCCCTTGCTctacgtggaagtggaagaagaggtgttaatcac |
| | | 795N.c5F | gtgattaacacctcttcttccacttccacgtagAGCAAGGGCGAGGAGCTGTTCACCGGGG |
| | | 795O.c2R | ttcataaaataaaatgtaggttacaattacCATtttgTATTgtcacttggttgttcacga |
| | | 795O.c3F | tcgtgaacaaccaagtgacAATAcaaaATGgtaattgtaacctacattttattttatgaa |
| | | 795O.c4R | CACCCCGGTGAACAGCTCCTCGCCCTTGCTctgtgggcacgatgatttttttatattagta |
| | | 795O.c5F | tactaatataaaaaatcatcgtgcccacagAGCAAGGGCGAGGAGCTGTTCACCGGGGTG |
| DsRed splicing PCR | | 795.s2F | CAATTGTGGCGTTTACAGCATTTGTTATACACACAGAACTC |
| | | 795.s2R | gaacagcatctgttacagcgacacaacatg |
| | | 795.s3F | CGATTCATCCTAGGctacaggaacaggtg |
| | | 795.s1R | catccaaccgccgccgcaacctgtc |
