## Supplementary Table 2 for "A fluorescent sex-sorting technique for insects with the demonstration in *Drosophila melanogaster*"

Supplementary Table 2: Sex-specific fluorescence sorting at multiple developmental stages.

| | Total embryos # | Total flies # | | | | | | | | | emerged adult | | | | | | emerged adult | | | | | | emerged adult | | |
|---|---|---|---|---|---|---|---|---|---|---|---|---|---|---|---|---|---|---|---|---|---|---|---|---|---|
| | | | egg# | larve# | hatching rate | total n+avg + std | pupa# | ess from adult su | avg + std | pupa R+G | ♀+♂ | ♀ | %♀ | ♂ | %♂ | pupa G | ♀+♂ | ♀ | %♀ | ♂ | %♂ | wt | ♀+♂ | ♀ | ♂ |
| w- | | | 160 | 156 | 97.5% | | 139 | 79.4% | | | | | | | | | | | | | | | 127 | 70 | 57 |
| w- | | | 159 | 156 | 98.1% | | 137 | 78.0% | | | | | | | | | | | | | | | 124 | 60 | 64 |
| w- | | | 203 | 194 | 95.6% | | 170 | 81.8% | | | | | | | | | | | | | | | 166 | 87 | 79 |
| w- | | | 202 | 200 | 99.0% | 97.11% | 178 | 86.1% | 82.68% | | | | | | | | | | | | | | 174 | 83 | 91 |
| w- | 918 | 762 | 194 | 185 | 95.4% | 1.60% | 179 | 88.1% | 4.35% | | | | | | | | | | | | | | 171 | 99 | 72 |
| 795G.1 | | | 168 | 128 | 76.2% | | 118 | 58.3% | | 118 | 98 | 52 | 53.1% | 46 | 46.94% | 0 | 0 | 0 | 0.00% | 0 | 0.00% | | | | |
| 795G.2 | | | 47 | 40 | 85.1% | | 34 | 59.6% | | 34 | 28 | 16 | 57.1% | 12 | 42.86% | 0 | 0 | 0 | 0.00% | 0 | 0.00% | | | | |
| 795G.3 | | | 111 | 79 | 71.2% | | 63 | 55.0% | | 63 | 61 | 30 | 49.2% | 31 | 50.82% | 0 | 0 | 0 | 0.00% | 0 | 0.00% | | | | |
| 795G.4 | | | 115 | 72 | 62.6% | 72.77% | 66 | 55.7% | 59.45% | 66 | 64 | 35 | 54.7% | 29 | 45.31% | 0 | 0 | 0 | 0.00% | 0 | 0.00% | | | | |
| 795G.5 | 521 | 306 | 80 | 55 | 68.8% | 8.45% | 55 | 68.8% | 5.53% | 55 | 55 | 28 | 50.9% | 27 | 49.09% | 0 | 0 | 0 | 0.00% | 0 | 0.00% | | | | |
| 795H.1 | | | 185 | 159 | 85.9% | | 108 | 35.1% | | 57 | 36 | 36 | 100.0% | 0 | 0.00% | 51 | 29 | 0 | 0.00% | 29 | 100.0% | | | | |
| 795H.2 | | | 175 | 113 | 64.6% | | 94 | 35.4% | | 47 | 34 | 34 | 100.0% | 0 | 0.00% | 47 | 28 | 0 | 0.00% | 28 | 100.0% | | | | |
| 795H.3 | | | 203 | 175 | 86.2% | | 156 | 17.7% | | 76 | 18 | 18 | 100.0% | 0 | 0.00% | 80 | 18 | 0 | 0.00% | 18 | 100.0% | | | | |
| 795H.4 | | | 232 | 193 | 83.2% | 79.49% | 145 | 52.6% | 39.61% | 67 | 59 | 59 | 100.0% | 0 | 0.00% | 78 | 63 | 0 | 0.00% | 63 | 100.0% | | | | |
| 795H.5 | 1040 | 425 | 245 | 190 | 77.6% | 9.04% | 167 | 57.1% | 15.75% | 86 | 79 | 79 | 100.0% | 0 | 0.00% | 81 | 61 | 0 | 0.00% | 61 | 100.0% | | | | |
| 795I.1 | | | 153 | 144 | 94.1% | | 143 | 56.9% | | 105 | 49 | 49 | 100.0% | 0 | 0.00% | 38 | 38 | 0 | 0.00% | 38 | 100.0% | | | | |
| 795I.2 | | | 216 | 191 | 88.4% | | 172 | 73.1% | | 92 | 87 | 87 | 100.0% | 0 | 0.00% | 80 | 71 | 0 | 0.00% | 71 | 100.0% | | | | |
| 795I.3 | | | 211 | 176 | 83.4% | | 156 | 60.2% | | 73 | 66 | 66 | 100.0% | 0 | 0.00% | 83 | 61 | 0 | 0.00% | 61 | 100.0% | | | | |
| 795I.4 | | | 290 | 227 | 78.3% | 82.64% | 225 | 62.4% | 64.63% | 89 | 84 | 84 | 100.0% | 0 | 0.00% | 136 | 97 | 0 | 0.00% | 97 | 100.0% | | | | |
| 795I.5 | 1136 | 720 | 266 | 214 | 80.5% | 4.39% | 175 | 62.8% | 5.79% | 88 | 83 | 83 | 100.0% | 0 | 0.00% | 87 | 84 | 0 | 0.00% | 84 | 100.0% | | | | |
| 795J.1 | | | 173 | 154 | 89.0% | | 152 | 74.0% | | 65 | 57 | 57 | 100.0% | 0 | 0.00% | 87 | 71 | 0 | 0.00% | 71 | 100.0% | | | | |
| 795J.2 | | | 148 | 141 | 95.3% | | 136 | 79.1% | | 57 | 53 | 53 | 100.0% | 0 | 0.00% | 79 | 64 | 0 | 0.00% | 64 | 100.0% | | | | |
| 795J.3 | | | 198 | 190 | 96.0% | | 159 | 70.7% | | 100 | 90 | 90 | 100.0% | 0 | 0.00% | 59 | 50 | 0 | 0.00% | 50 | 100.0% | | | | |
| 795J.4 | | | 212 | 201 | 94.8% | 94.32% | 168 | 66.0% | 71.52% | 86 | 74 | 74 | 100.0% | 0 | 0.00% | 82 | 66 | 0 | 0.00% | 66 | 100.0% | | | | |
| 795J.5 | 933 | 662 | 202 | 195 | 96.5% | 3.04% | 164 | 67.8% | 5.18% | 76 | 69 | 69 | 100.0% | 0 | 0.00% | 88 | 68 | 0 | 0.00% | 68 | 100.0% | | | | |
| 795K.1 | | | 256 | 224 | 87.5% | | 205 | 75.8% | | 205 | 194 | 102 | 52.6% | 92 | 47.42% | 0 | 0 | 0 | 0.00% | 0 | 0.00% | | | | |
| 795K.2 | | | 212 | 180 | 84.9% | | 172 | 76.9% | | 172 | 163 | 79 | 48.5% | 84 | 51.53% | 0 | 0 | 0 | 0.00% | 0 | 0.00% | | | | |
| 795K.3 | | | 188 | 180 | 95.7% | | 125 | 57.4% | | 125 | 108 | 61 | 56.5% | 47 | 43.52% | 0 | 0 | 0 | 0.00% | 0 | 0.00% | | | | |
| 795K.4 | | | 207 | 172 | 83.1% | 87.68% | 160 | 69.6% | 72.73% | 160 | 144 | 80 | 55.6% | 64 | 44.44% | 0 | 0 | 0 | 0.00% | 0 | 0.00% | | | | |
| 795K.5 | 1050 | 766 | 187 | 163 | 87.2% | 4.85% | 159 | 84.0% | 9.95% | 159 | 157 | 88 | 56.1% | 69 | 43.95% | 0 | 0 | 0 | 0.00% | 0 | 0.00% | | | | |
| | | | egg# | larve# | hatching rate | avg + std | pupa# | ess from adult su | avg + std | pupa R+G | ♀+♂ | ♀ | %♀ | ♂ | %♂ | pupa R | ♀+♂ | ♀ | %♀ | ♂ | %♂ | | | | |
| 795L.1 | | | 393 | 254 | 64.6% | | 235 | 53.4% | | 120 | 108 | 108 | 100.0% | 0 | 0.00% | 115 | 102 | 0 | 0.00% | 102 | 100.0% | | | | |
| 795L.2 | | | 214 | 173 | 80.8% | | 160 | 62.1% | | 70 | 62 | 62 | 100.0% | 0 | 0.00% | 90 | 71 | 0 | 0.00% | 71 | 100.0% | | | | |
| 795L.3 | | | 244 | 217 | 88.9% | | 173 | 63.9% | | 75 | 69 | 69 | 100.0% | 0 | 0.00% | 98 | 87 | 0 | 0.00% | 87 | 100.0% | | | | |
| 795L.4 | | | 209 | 157 | 75.1% | 77.10% | 127 | 54.1% | 57.89% | 65 | 57 | 57 | 100.0% | 0 | 0.00% | 62 | 56 | 0 | 0.00% | 56 | 100.0% | | | | |
| 795L.5 | 1239 | 712 | 179 | 136 | 76.0% | 8.87% | 113 | 55.9% | 4.83% | 63 | 55 | 55 | 100.0% | 0 | 0.00% | 50 | 45 | 0 | 0.00% | 45 | 100.0% | | | | |
| 795M.1 | | | 102 | 83 | 81.4% | | 61 | 52.94% | | 34 | 29 | 29 | 100.0% | 0 | 0.00% | 27 | 25 | 0 | 0.00% | 25 | 100.0% | | | | |
| 795M.2 | | | 63 | 46 | 73.0% | | 46 | 65.08% | | 23 | 20 | 20 | 100.0% | 0 | 0.00% | 23 | 21 | 0 | 0.00% | 21 | 100.0% | | | | |
| 795M.3 | | | 94 | 68 | 72.3% | | 44 | 45.74% | | 25 | 24 | 24 | 100.0% | 0 | 0.00% | 19 | 19 | 0 | 0.00% | 19 | 100.0% | | | | |
| 795M.4 | | | 132 | 118 | 89.4% | 76.62% | 54 | 37.88% | 49.87% | 29 | 25 | 25 | 100.0% | 0 | 0.00% | 25 | 25 | 0 | 0.00% | 25 | 100.0% | | | | |
| 795M.5 | 500 | 240 | 109 | 73 | 67.0% | 8.81% | 53 | 47.71% | 10.08% | 31 | 31 | 31 | 100.0% | 0 | 0.00% | 22 | 21 | 0 | 0.00% | 21 | 100.0% | | | | |
| 795N.1 | | | 181 | 147 | 81.2% | | 132 | 58.56% | | 70 | 61 | 61 | 100.0% | 0 | 0.00% | 62 | 45 | 0 | 0.00% | 45 | 100.0% | | | | |
| 795N.2 | | | 197 | 167 | 84.8% | | 117 | 49.24% | | 74 | 55 | 55 | 100.0% | 0 | 0.00% | 43 | 42 | 0 | 0.00% | 42 | 100.0% | | | | |
| 795N.3 | | | 435 | 296 | 68.0% | | 229 | 45.06% | | 125 | 114 | 114 | 100.0% | 0 | 0.00% | 104 | 82 | 0 | 0.00% | 82 | 100.0% | | | | |
| 795N.4 | | | 186 | 158 | 84.9% | 76.89% | 105 | 46.24% | 48.76% | 56 | 42 | 42 | 100.0% | 0 | 0.00% | 49 | 44 | 0 | 0.00% | 44 | 100.0% | | | | |
| 795N.5 | 1225 | 586 | 226 | 148 | 65.5% | 9.41% | 127 | 44.69% | 5.77% | 68 | 48 | 48 | 100.0% | 0 | 0.00% | 59 | 53 | 0 | 0.00% | 53 | 100.0% | | | | |
| 795O.1 | | | 233 | 169 | 72.5% | | 128 | 49.79% | | 128 | 116 | 60 | 51.7% | 56 | 48.28% | 0 | 0 | 0 | 0.00% | 0 | 0.0% | | | | |
| 795O.2 | | | 280 | 209 | 74.6% | | 106 | 35.71% | | 106 | 100 | 50 | 50.0% | 50 | 50.00% | 0 | 0 | 0 | 0.00% | 0 | 0.0% | | | | |
| 795O.3 | | | 214 | 165 | 77.1% | | 114 | 50.00% | | 114 | 107 | 57 | 53.3% | 50 | 46.73% | 0 | 0 | 0 | 0.00% | 0 | 0.0% | | | | |
| 795O.4 | | | 188 | 134 | 71.3% | 73.96% | 107 | 55.32% | 50.27% | 107 | 104 | 54 | 51.9% | 50 | 48.08% | 0 | 0 | 0 | 0.00% | 0 | 0.0% | | | | |
| 795O.5 | 1148 | 568 | 233 | 173 | 74.2% | 2.22% | 154 | 60.52% | 9.26% | 154 | 141 | 63 | 44.7% | 78 | 55.32% | 0 | 0 | 0 | 0.00% | 0 | 0.0% | | | | |
| | | | | | | | | | | | emerged adult | | | | | | emerged adult | | | | | | | | |
| | | | egg# | L1# | hatching rate | avg + std | L3# | L3 survival rate | avg + std | L3 R+G | ♀+♂ | ♀ | %♀ | ♂ | %♂ | L3 G | ♀+♂ | ♀ | %♀ | ♂ | %♂ | | | | |
| 795H.1 | | | 218 | 172 | 78.90% | | 152 | 69.72% | | 74 | 51 | 51 | 100.00% | 0 | 0.00% | 78 | 57 | 0 | 0.00% | 57 | 100.00% | | | | |
| 795H.2 | | | 230 | 166 | 72.17% | | 138 | 60.00% | | 65 | 51 | 51 | 100.00% | 0 | 0.00% | 73 | 53 | 0 | 0.00% | 53 | 100.00% | | | | |
| 795H.3 | | | 239 | 213 | 89.12% | | 156 | 65.27% | | 76 | 54 | 54 | 100.00% | 0 | 0.00% | 80 | 64 | 0 | 0.00% | 64 | 100.00% | | | | |
| 795H.4 | | | 215 | 197 | 91.63% | | 146 | 67.91% | 65.39% | 88 | 52 | 52 | 100.00% | 0 | 0.00% | 58 | 49 | 0 | 0.00% | 49 | 100.00% | | | | |
| 795H.5 | 1130 | 553 | 228 | 188 | 82.46% | | 146 | 64.04% | 3.74% | 76 | 53 | 53 | 100.00% | 0 | 0.00% | 70 | 69 | 0 | 0.00% | 69 | 100.00% | | | | |
| 795I.1 | | | 97 | 94 | 96.91% | | 86 | 88.66% | | 41 | 41 | 41 | 100.00% | 0 | 0.00% | 45 | 44 | 0 | 0.00% | 44 | 100.00% | | | | |
| 795I.2 | | | 95 | 87 | 91.58% | | 87 | 91.58% | | 45 | 41 | 41 | 100.00% | 0 | 0.00% | 42 | 29 | 0 | 0.00% | 29 | 100.00% | | | | |
| 795I.3 | | | 225 | 196 | 87.11% | | 185 | 82.22% | | 102 | 92 | 92 | 100.00% | 0 | 0.00% | 83 | 74 | 0 | 0.00% | 74 | 100.00% | | | | |
| 795I.4 | | | 210 | 195 | 92.86% | 89.50% | 181 | 86.19% | 80.99% | 93 | 85 | 85 | 100.00% | 0 | 0.00% | 88 | 79 | 0 | 0.00% | 79 | 100.00% | | | | |
| 795I.5 | 842 | 596 | 215 | 170 | 79.07% | 6.80% | 121 | 56.28% | 14.23% | 58 | 57 | 57 | 100.00% | 0 | 0.00% | 63 | 54 | 0 | 0.00% | 54 | 100.00% | | | | |
| 795J.1 | | | 191 | 175 | 91.62% | | 170 | 89.01% | | 94 | 86 | 86 | 100.00% | 0 | 0.00% | 76 | 71 | 0 | 0.00% | 71 | 100.00% | | | | |
| 795J.2 | | | 251 | 226 | 90.04% | | 220 | 87.65% | | 120 | 106 | 106 | 100.00% | 0 | 0.00% | 100 | 75 | 0 | 0.00% | 75 | 100.00% | | | | |
| 795J.3 | | | 159 | 155 | 97.48% | | 146 | 91.82% | | 86 | 72 | 72 | 100.00% | 0 | 0.00% | 60 | 54 | 0 | 0.00% | 54 | 100.00% | | | | |
| 795J.4 | | | 226 | 184 | 81.42% | 90.39% | 180 | 79.65% | 86.81% | 91 | 77 | 77 | 100.00% | 0 | 0.00% | 89 | 80 | 0 | 0.00% | 80 | 100.00% | | | | |
| 795J.5 | 1118 | 838 | 291 | 266 | 91.41% | 5.78% | 250 | 85.91% | 4.55% | 140 | 132 | 132 | 100.00% | 0 | 0.00% | 110 | 85 | 0 | 0.00% | 85 | 100.00% | | | | |
| | | | egg# | L1# | hatching rate | avg + std | L3# | L3 survival rate | avg + std | L3 R+G | ♀+♂ | ♀ | %♀ | ♂ | %♂ | L3 R | ♀+♂ | ♀ | %♀ | ♂ | %♂ | | | | |
| 795L.1 | | | 193 | 167 | 86.53% | | 161 | 83.42% | | 72 | 49 | 49 | 100.00% | 0 | 0.00% | 89 | 42 | 0 | 0.00% | 42 | 100.00% | | | | |
| 795L.2 | | | 222 | 177 | 79.73% | | 145 | 65.32% | | 69 | 30 | 30 | 100.00% | 0 | 0.00% | 76 | 34 | 0 | 0.00% | 34 | 100.00% | | | | |
| 795L.3 | | | 131 | 110 | 83.97% | | 81 | 61.83% | | 26 | 15 | 15 | 100.00% | 0 | 0.00% | 55 | 25 | 0 | 0.00% | 25 | 100.00% | | | | |
| 795L.4 | | | 134 | 110 | 82.09% | 82.15% | 84 | 62.69% | 65.49% | 47 | 30 | 30 | 100.00% | 0 | 0.00% | 37 | 15 | 0 | 0.00% | 15 | 100.00% | | | | |
| 795L.5 | 907 | 313 | 227 | 188 | 82.82% | 1.79% | 123 | 54.19% | 10.85% | 71 | 48 | 48 | 100.00% | 0 | 0.00% | 52 | 25 | 0 | 0.00% | 25 | 100.00% | | | | |
| | | | | | | | | | | | emerged adult | | | | | | emerged adult | | | | | | | | |
| | | | egg# | L1# | hatching rate | avg + std | L2# | L2 survival rate | avg + std | L2 R+G | ♀+♂ | ♀ | %♀ | ♂ | %♂ | L2 G | ♀+♂ | ♀ | %♀ | ♂ | %♂ | | | | |
| 795H.1 | | | 252 | 205 | 81.35% | | 158 | 62.70% | | 81 | 72 | 72 | 100.00% | 0 | 0.00% | 77 | 73 | 0 | 0.00% | 73 | 100.00% | | | | |
| 795H.2 | | | 237 | 197 | 83.12% | | 185 | 78.06% | | 94 | 90 | 90 | 100.00% | 0 | 0.00% | 91 | 84 | 0 | 0.00% | 84 | 100.00% | | | | |
| 795H.3 | | | 195 | 161 | 82.56% | | 140 | 71.79% | | 66 | 64 | 64 | 100.00% | 0 | 0.00% | 74 | 68 | 0 | 0.00% | 68 | 100.00% | | | | |
| 795H.4 | | | 211 | 170 | 80.57% | 79.91% | 116 | 54.98% | 66.49% | 53 | 41 | 41 | 100.00% | 0 | 0.00% | 63 | 52 | 0 | 0.00% | 52 | 100.00% | | | | |
| 795H.5 | 1123 | 686 | 228 | 164 | 71.93% | 4.57% | 148 | 64.91% | 8.82% | 78 | 72 | 72 | 100.00% | 0 | 0.00% | 70 | 70 | 0 | 0.00% | 70 | 100.00% | | | | |
| 795I.1 | | | 161 | 144 | 89.44% | | 89 | 55.28% | | 47 | 46 | 46 | 100.00% | 0 | 0.00% | 42 | 33 | 0 | 0.00% | 33 | 100.00% | | | | |
| 795I.2 | | | 108 | 99 | 91.67% | | 95 | 87.96% | | 46 | 46 | 46 | 100.00% | 0 | 0.00% | 49 | 48 | 0 | 0.00% | 48 | 100.00% | | | | |
| 795I.3 | | | 136 | 121 | 88.97% | | 110 | 80.88% | | 59 | 58 | 58 | 100.00% | 0 | 0.00% | 51 | 51 | 0 | 0.00% | 51 | 100.00% | | | | |
| 795I.4 | | | 73 | 65 | 89.04% | 89.97% | 62 | 84.93% | 75.51% | 32 | 21 | 21 | 100.00% | 0 | 0.00% | 30 | 28 | 0 | 0.00% | 28 | 100.00% | | | | |
| 795I.5 | 532 | 363 | 54 | 49 | 90.74% | 1.18% | 37 | 68.52% | 13.52% | 20 | 19 | 19 | 100.00% | 0 | 0.00% | 17 | 13 | 0 | 0.00% | 13 | 100.00% | | | | |
| | | | egg# | L1# | hatching rate | avg + std | L2# | L2 survival rate | avg + std | L2 R+G | ♀+♂ | ♀ | %♀ | ♂ | %♂ | L3 R | ♀+♂ | ♀ | %♀ | ♂ | %♂ | | | | |
| 795L.1 | | | 227 | 205 | 90.31% | | 199 | 87.67% | | 97 | 66 | 66 | 100.00% | 0 | 0.00% | 102 | 42 | 0 | 0.00% | 42 | 100.00% | | | | |
| 795L.2 | | | 96 | 63 | 65.63% | | 54 | 56.25% | | 25 | 20 | 20 | 100.00% | 0 | 0.00% | 29 | 18 | 0 | 0.00% | 18 | 100.00% | | | | |
| 795L.3 | | | 92 | 55 | 59.78% | | 47 | 51.09% | | 27 | 18 | 18 | 100.00% | 0 | 0.00% | 20 | 8 | 0 | 0.00% | 8 | 100.00% | | | | |
| 795L.4 | | | 135 | 102 | 75.56% | 72.09% | 80 | 59.26% | 60.76% | 43 | 24 | 24 | 100.00% | 0 | 0.00% | 37 | 20 | 0 | 0.00% | 20 | 100.00% | | | | |
| 795L.5 | 657 | 262 | 107 | 74 | 69.16% | 11.68% | 53 | 49.53% | 15.54% | 27 | 22 | 22 | 100.00% | 0 | 0.00% | 26 | 24 | 0 | 0.00% | 24 | 100.00% | | | | |
| | | | | | | | | | | | | | | | emerged adult | | | emerged adult | | | | | | | |
| | | | egg# | L1# | hatching rate | avg + std | L1 R+G | ♀+♂ | ♀ | %♀ | ♂ | %♂ | L2 G | ♀+♂ | ♀ | %♀ | ♂ | %♂ | | | | | | | |
| 795H.1 | | | 89 | 86 | 96.63% | | 41 | 26 | 26 | 100.00% | 0 | 0.00% | 45 | 30 | 0 | 0.00% | 30 | 100.00% | | | | | | | |
| 795H.2 | | | 160 | 157 | 98.13% | | 74 | 50 | 50 | 100.00% | 0 | 0.00% | 83 | 36 | 0 | 0.00% | 36 | 100.00% | | | | | | | |
| 795H.3 | | | 143 | 139 | 97.20% | | 66 | 47 | 47 | 100.00% | 0 | 0.00% | 73 | 34 | 0 | 0.00% | 34 | 100.00% | | | | | | | |
| 795H.4 | | | 128 | 126 | 98.44% | 97.40% | 67 | 40 | 40 | 100.00% | 0 | 0.00% | 59 | 39 | 0 | 0.00% | 39 | 100.00% | | | | | | | |
| 795H.5 | 637 | 362 | 117 | 113 | 96.58% | 0.85% | 61 | 31 | 31 | 100.00% | 0 | 0.00% | 52 | 29 | 0 | 0.00% | 29 | 100.00% | | | | | | | |
| | | | egg# | L1# | hatching rate | avg + std | L1 R+G | ♀+♂ | ♀ | %♀ | ♂ | %♂ | L2 R | ♀+♂ | ♀ | %♀ | ♂ | %♂ | | | | | | | |
| 795L.1 | | | 106 | 91 | 85.85% | | 45 | 38 | 38 | 100.00% | 0 | 0.00% | 46 | 38 | 0 | 0.00% | 38 | 100.00% | | | | | | | |
| 795L.2 | | | 133 | 102 | 76.69% | | 53 | 45 | 45 | 100.00% | 0 | 0.00% | 49 | 27 | 0 | 0.00% | 27 | 100.00% | | | | | | | |
| 795L.3 | | | 145 | 109 | 75.17% | | 61 | 45 | 45 | 100.00% | 0 | 0.00% | 48 | 39 | 0 | 0.00% | 39 | 100.00% | | | | | | | |
| 795L.4 | | | 145 | 124 | 85.52% | 80.48% | 65 | 50 | 50 | 100.00% | 0 | 0.00% | 59 | 39 | 0 | 0.00% | 39 | 100.00% | | | | | | | |
| 795L.5 | 673 | 427 | 144 | 114 | 79.17% | 4.96% | 62 | 62 | 62 | 100.00% | 0 | 0.00% | 52 | 44 | 0 | 0.00% | 44 | 100.00% | | | | | | | |
